# Deep learning-based algorithm for predicting the live birth potential of mouse embryos

**DOI:** 10.1101/2021.08.19.456065

**Authors:** Yuta Tokuoka, Takahiro G. Yamada, Daisuke Mashiko, Zenki Ikeda, Tetsuya J. Kobayashi, Kazuo Yamagata, Akira Funahashi

**Affiliations:** Center for Biosciences and Informatics, Graduate School of Fundamental Science and Technology, Keio University, Kanagawa, Japan; Department of Biosciences and Informatics, Keio University, Kanagawa, Japan; Faculty of Biology-Oriented Science and Technology, Kindai University, Wakayama, Japan; Institute of Industrial Science, The University of Tokyo, Tokyo, Japan

## Abstract

In assisted reproductive technology (ART), embryos produced by *in vitro* fertilization (IVF) are graded according to their live birth potential, and high-grade embryos are preferentially transplanted. However, the rate of live birth following clinical ART remains low worldwide, suggesting that grading is inaccurate. One explanation is that grading is classically based on the characteristic shape of embryos at a limited number of developmental stages and does not consider the shape of embryos and intracellular structures, e.g., nuclei, at various stages important for normal embryogenesis. Therefore, here we developed a Normalized Multi-View Attention Network (NVAN) that directly predicts live birth potential from nuclear structural features in live-cell fluorescence images taken of mouse embryos across a wide range of stages. The classification accuracy of our method was 83.87%, which greatly exceeded that of existing machine-learning methods and that of visual inspection by embryo culture specialists. By visualizing the features that contributed most to the prediction of live birth potential, we found that the size and shape of the cell nucleus at the morula stage and at the time of cell division were important for live birth prediction. We anticipate that our method will help ART and developmental engineering as a new basic technology for IVF embryo selection.

## Introduction

In the field of developmental engineering, various mammalian embryos have been produced by IVF, cultured *in vitro*, and transplanted to foster mothers, and this series of technologies are called assisted reproductive technologies (ART). For example, human, bovine, and mouse embryos have been used for infertility treatment [1, 2], livestock production [3, 4], and genetic modification and genome editing technologies [5, 6], respectively.

During ART, embryo grading based on embryo morphology is commonly used to select embryos with a high potential of live birth: e.g., grading using Veeck criteria, International Embryo Technology Society (IETS) criteria, and Gardner criteria [7–11]. High-grade embryos are then preferentially transplanted into the mother for the purpose of increasing the birth rate. However, since the embryologist manually and subjectively grades the embryos, the grading results vary among embryologists [2, 12] and the time cost of grading is enormous [13]. Therefore, in recent years, in an attempt to unify the evaluation of embryo grade, several studies have reported the application of machine learning to microscopic images for the grading of embryos [1, 4, 14, 15]. Rocha et al. trained Artificial Neural Networks (ANNs) on microscopic images of bovine embryos graded by IETS criteria [4]. From the images at day 7 post-fertilization, their trained ANN was able to predict embryo grade with 76.4% accuracy. Khosravi et al. trained Convolutional Neural Networks (CNNs) on microscopic images of human embryos graded by Veeck criteria [1]. From images at a single time point randomly selected in the developmental process, their trained CNN was able to predict embryo grade with 96.94% accuracy; however, the prediction accuracy of live birth using a similar CNN was only 51.85% [1]. Although the use of embryo grading has improved the live birth rate of human IVF embryos in ART worldwide, the rate remains at 28.0% [16]. Therefore, there is room for improvement in the grading of embryos for embryo selection.

Traditionally, embryo grading is based on the condition of the embryo during limited stages of embryogenesis. For example, the Veeck criteria are assessed at a narrow period within the early stages of development (between 4 and 16 cell stage) [11], and the IETS criteria and Gardner criteria are assessed at a specific part of the blastocyst stage (between 32 and 64 cell stage) [8, 10]. However, it is known that embryonic morphology changes dynamically and profoundly over time during early embryogenesis [17], and that embryos that develop into healthy blastocysts need to have a uniform time interval from one cell division to the next during the very early stages of development (between 1 and 4 cell stage) [18]. Because time is clearly such an important factor in early development, we expected that highly accurate prediction of embryos with live birth potential could be achieved by considering the time at various stages in early development.

Existing embryo evaluation methods using machine learning have focused only on improving prediction accuracy, and improving the interpretability of prediction outcomes has been largely ignored. On the other hand, in recent years, attempts to improve the interpretability of prediction outcomes have been made in other medical fields such as prostate cancer prognostics [19]. Low explainability of the predictions made by machine learning algorithms not only makes it difficult to assess the reliability of these predictions in clinical practice but is also a lost opportunity for physician decision-making and patient-centred care [20]. Therefore, a highly explainable embryo evaluation method would be indispensable in clinical practice and in ART generally.

In this study, we propose a deep learning-based method that can (1) predict the live birth potential of mouse embryos by considering the nuclear features of the embryos at various stages during early development and (2) explain the basis of the prediction outcomes (Fig. 1). In our method, for each embryo, the input is time-series microscopic fluorescent images of nuclear dynamics during early cleavage and the output is the binary classification of live or not live birth. Our method used Quantitative Criteria Acquisition Network (QCANet) [17], which can perform segmentation of cell nuclei in microscopic images, to extract multivariate time-series data of embryo behaviour (including the number, shape, and position of cell nuclei in a time series) at each time point; among existing methods, this method enables cell nucleus segmentation of mouse early embryos with the highest accuracy [17]. We developed Normalized Multi-View Attention Network (NVAN), which uses a novel attention mechanism, to classify embryos from multivariate time-series data. With NVAN, we were able to accurately predict the live birth potential of the embryos on the basis of nuclear features at various stages during early development. Furthermore, by visualizing the features that contributed to the prediction of live birth potential by NVAN, we found that the size and shape of the cell nuclei at the time of the cell divisions leading to the morula stage and at the morula were important for birth prediction. Further analysis of these features suggested that conditions such as variation in the size and shape of cell nuclei were necessary for the prediction of live birth potential. This variation in size and shape of cell nuclei is a promising criterion for embryo grading to improve the live birth rate.

**Figure 1:**
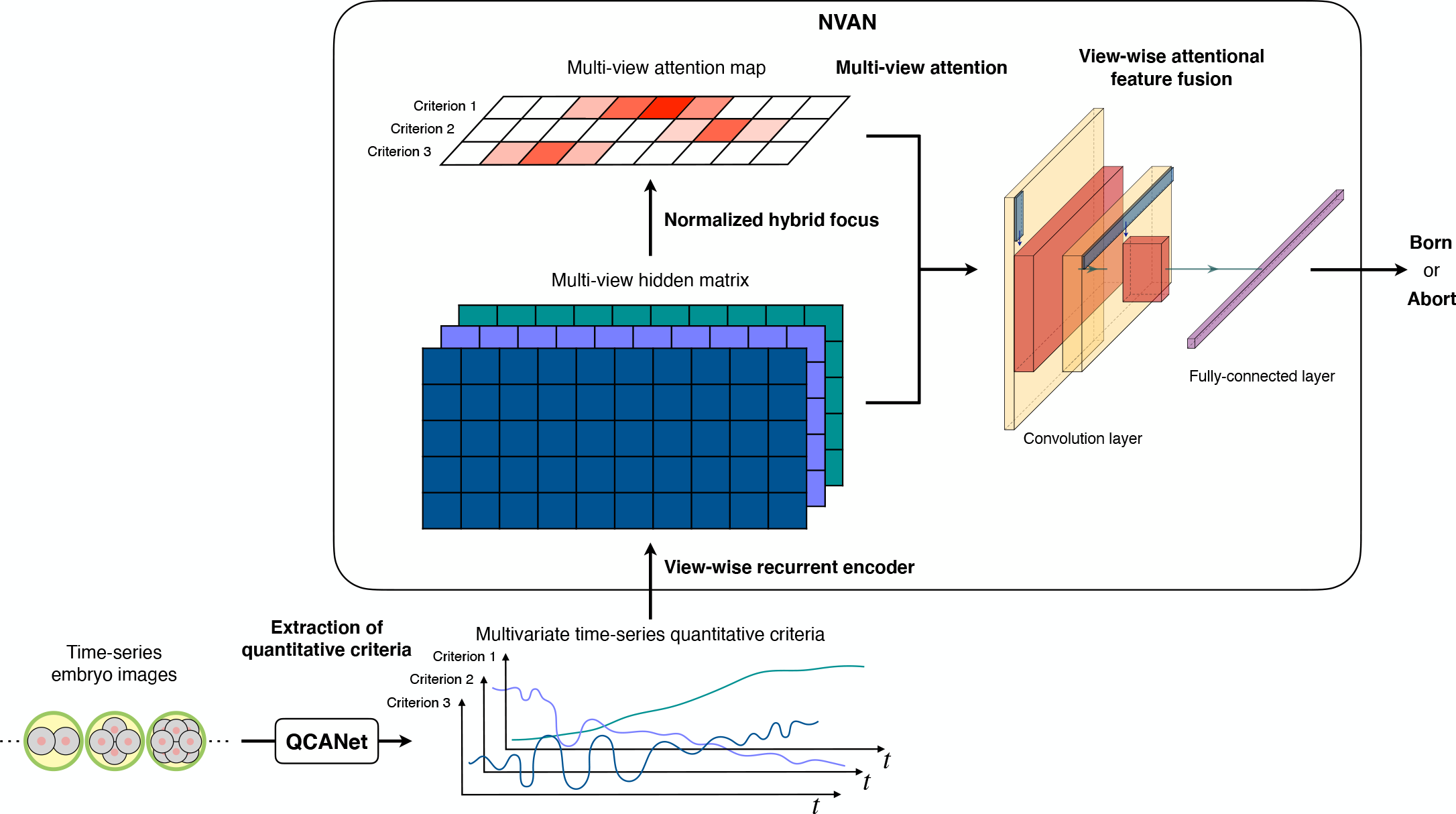
Conceptual diagram of our method. We first used Quantitative Criteria Acquisition Network (QCANet) [17] to perform segmentation of a published time series of 3D fluorescence microscopy images acquired by live cell imaging [21]. The resultant multivariate time-series data describing embryo behaviour were then used as input for a classification algorithm, Normalized Multi-View Attention Network (NVAN), to predict whether the embryo is live born or not (“born”/”abort”) (see Methods).

## Results

### Extraction of nuclei-based quantitative parameters for embryo development

We used published live-cell imaging data of early mouse embryos from a single blastocyst transfer experiment [21] and additional data that was obtained under the same experimental conditions as that used in [21]. The dataset consists of time-series three-dimensional (3D) fluorescence microscopic images (2,359,200 images) of 91 early mouse embryos that were labelled with a fluorescent protein-labelled histone, histone H2B–mCherry, for approximately 3.5 days from immediately after fertilization to the blastocyst stage. In the dataset each embryo is labelled as “born” or “abort” according to whether the blastocyst transfer resulted in a live birth or not, respectively: there were 62 born embryos and 29 abort embryos.

By using QCANet, we performed segmentation for cell nuclei of all embryos in the single blastocyst transfer dataset. Time-series images after 2.5 days post-fertilization are reported to display reduced segmentation accuracy compared with earlier images [21], and they were therefore excluded from the dataset. From the remaining images, 11 variables relating to embryo behaviour were extracted (Fig. 2): number of cell nuclei (number), mean volume of cell nuclei (volume_mean), standard deviation of cell nuclei volume (volume_sd), mean surface area of cell nuclei (surface_mean), standard deviation of cell nuclei surface area (surface_sd), mean aspect ratio (major axis/minor axis) of cell nuclei (aspect_ratio_mean), standard deviation of cell nuclei aspect ratio (aspect_ratio_sd), mean solidity (ratio of total area of the nucleus to the area of the convex hull) of cell nuclei (solidity_mean), standard deviation of cell nuclei solidity (solidity_sd), mean distance between embryo centre and every cell nucleus (centroid_mean), and standard deviation of the distance between embryo centre and every cell nucleus (centroid_sd).

**Figure 2:**
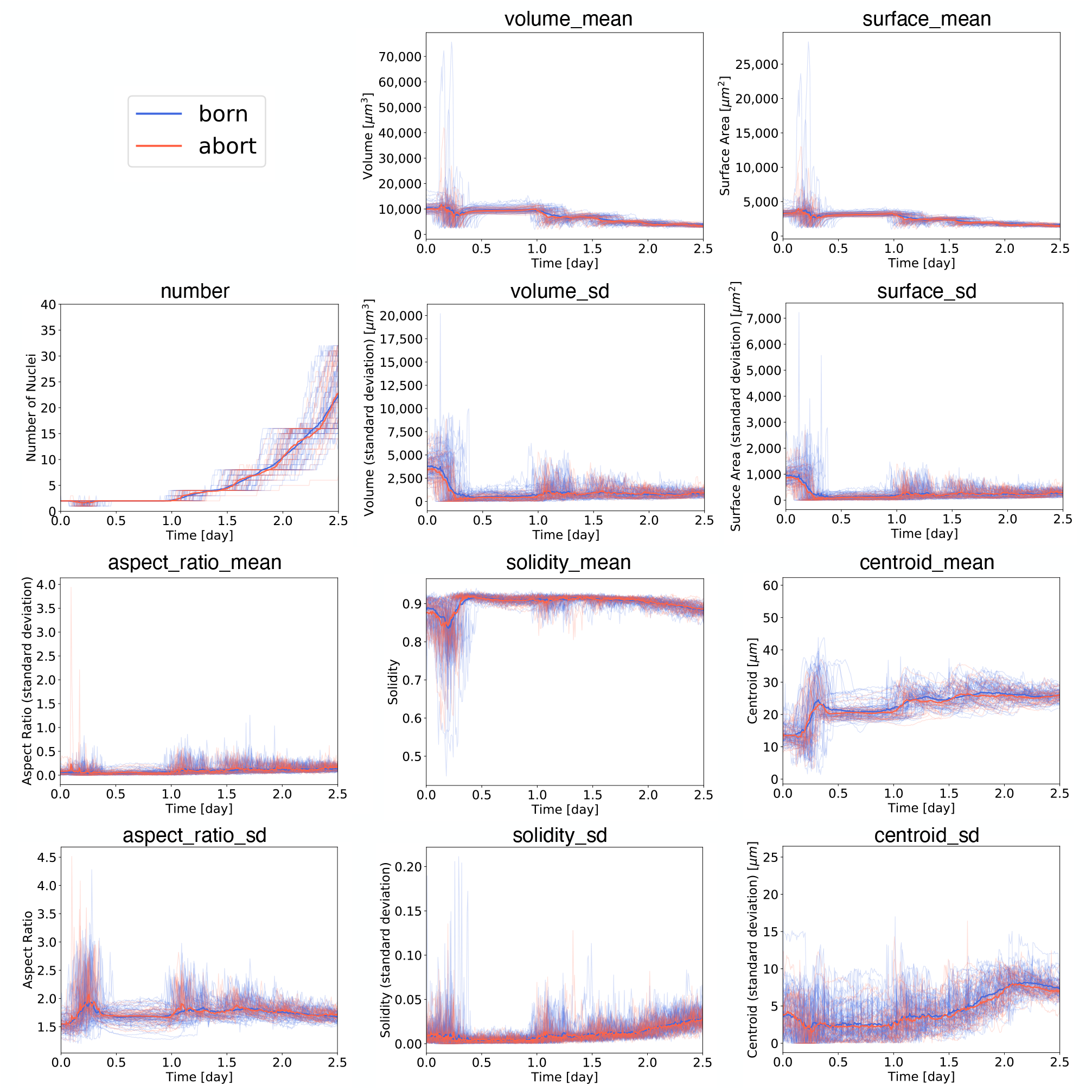
Time-series data of 11 criteria during the development of each embryo. Blue and red series with thin lines represent the values of quantitative criteria for individual embryos in the born and abort groups, respectively. Blue and red series with bold lines represent the means of the born or abort embryo groups, respectively. Volume_mean, mean volume of cell nuclei; surface_mean, mean surface area of cell nuclei; number, number of cell nuclei; volume_sd, standard deviation of cell nuclei volume; surface_sd, standard deviation of cell nuclei surface area; aspect_ratio_mean, mean aspect ratio of cell nuclei; solidity_mean, mean solidity of cell nuclei; centroid_mean, mean distance between embryo centre and every cell nucleus; aspect_ratio_sd, standard deviation of cell nuclei aspect ratio; solidity_sd, standard deviation of cell nuclei solidity; and centroid_sd, standard deviation of the distance between embryo centre and every cell nucleus.

When we analysed each of the 11 variables separately, no significant difference was found between the born and abort groups according to the two-sided Wilcoxon rank-sum test (Benjamini-Hochberg adjusted *p*-value *>* 0.05). From this result, we conclude that it is diffcult to comprehensively classify the live birth potential of embryos on the basis of the behaviour of a single variable. To circumvent this issue, we therefore developed a new classification method based on the multivariate time-series data, as described below.

### Accuracy comparison for prediction of live birth potential by our method versus existing machine-learning methods

We used the data acquired for the 11 variables of embryo behaviour as multivariate time-series data to train a novel algorithm for the prediction of an embryo’s live birth potential (i.e., born/abort classification). The proposed algorithm, NVAN, is based on Long Short-Term Memory (LSTM) [22] (see Methods). To evaluate how superior NVAN is for classifying time-series data, we trained and evaluated other neural network algorithms used for classifying time-series data, namely LSTM [22], AttentionLSTM [23], and Transformer [24], as well as the highly accurate MuVAN [25], TG-LSTM [26], and TapNet [27]. We also trained and evaluated XGBoost [28], a decision-tree algorithm, because decision trees are famous for their high interpretability. All of these algorithms, like NVAN, were trained by using the multivariate time-series data as input. In addition, we trained and evaluated CLDNN [29], which is an algorithm for predicting the live birth potential of embryos directly from time-series images: i.e., without using the multivariate time-series data. Because the images in the dataset include time-series two-dimensional (2D) bright-field microscopic images and time-series 3D fluorescence microscopic images, we prepared three patterns of learning machine: 2DCLDNN (input, 2D images), 3DCLDNN (input, 3D images), and 5DCLDNN (input, 5D images that concatenated 2D images and 3D images).

To evaluate the classification accuracy of each method, we randomly divided the entire dataset into training and test sets at a ratio of 2 : 1. The training data were further repeatedly and randomly divided into training and validation sets at a ratio of 3 : 1 in 4-fold cross-validation, and the most accurate model for each method was determined. The best model for each method was then evaluated on the test data, and the classification accuracy of each method was calculated. We used Accuracy (correct answer rate), F-measure (comprehensive measure of absence of false negatives and false positives), AUROC (area under the receiver operating characteristic), and AUPR (area under the precision-recall) to evaluate the robustness of the classification accuracy of the learning model. The classification accuracy and robustness of our method using NVAN and the multivariate time-series data exceeded that of all the other methods in all the evaluation metrics (Accuracy, 0.8387; F-measure, 0.8980; AUROC, 0.7879; AUPR, 0.8855) (Fig. 3, Supplementary Table 1; Supplementary Fig. 1).

**Figure 3:**
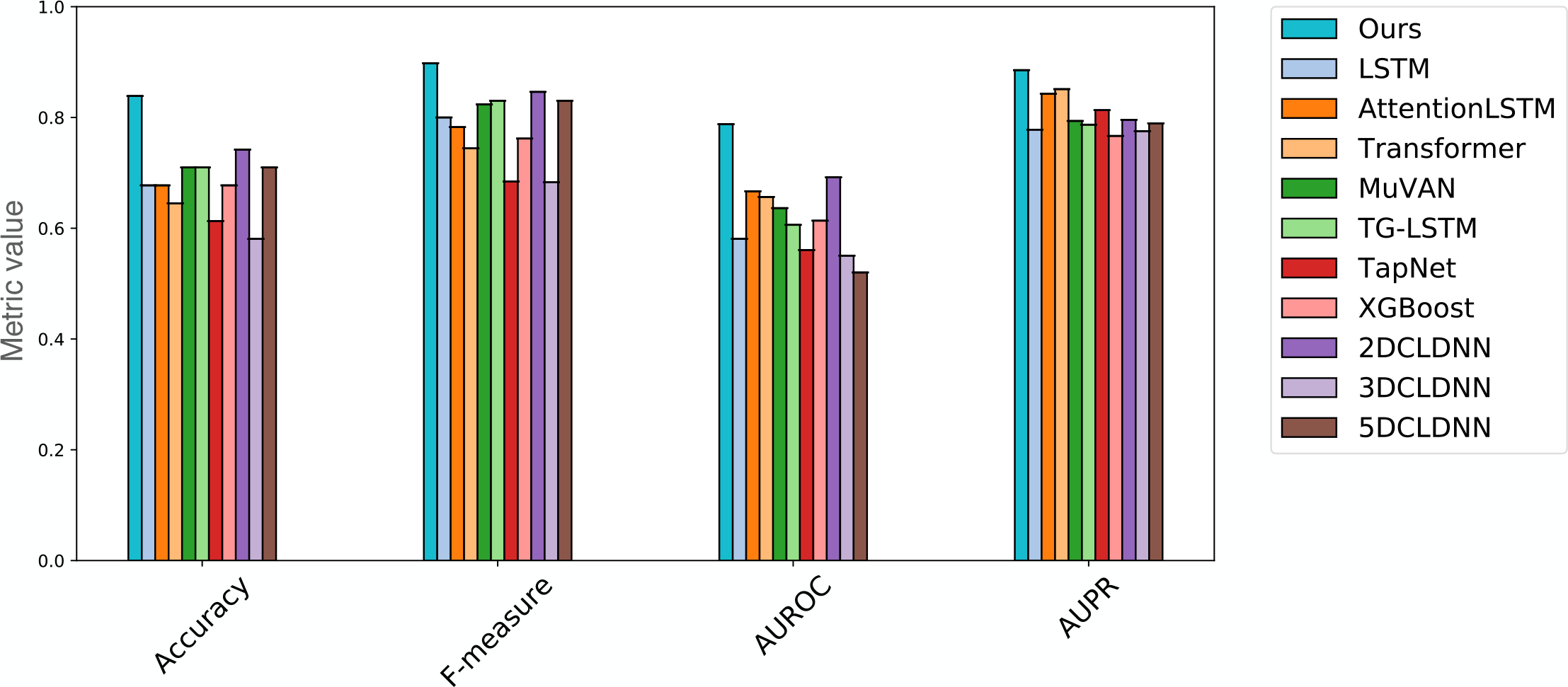
Comparison of the classification accuracy of each method. We compared the accuracy of our method (Ours), with those of LSTM [22], AttentionLSTM [23], Transformer [24], MuVAN [25], TG-LSTM [26], TapNet [27], XGBoost [28], 2DCLDNN [29], 3DCLDNN [29], and 5DCLDNN [29]. The classification accuracy of each method is shown for various classification metrics (Accuracy, F-measure, AUROC, and AUPR).

### Comparison of classification accuracy for prediction of live birth potential between our method and embryo culture specialists

When embryo culture specialists make a morphological evaluation of embryos, they grade the embryos on the basis of the shape of the developing embryos observed under a bright-field microscope. Here, nine embryo culture specialists observed time-series bright-field microscopic images of the embryos used to create the above test data and then classified each embryo as born or abort. We then compared their classification accuracy to that of our method (Fig. 4, Supplementary Table 2). The classification accuracy of our method (Accuracy 0.8387, F-measure 0.8980) greatly exceeded that of the embryo culture specialists (Accuracy 0.6422, F-measure 0.7314). This result shows that the predictions made by our method not only exceeded the classification accuracy of existing machine-learning methods but also exceeded the accuracy of embryo selection by specialists.

**Figure 4:**
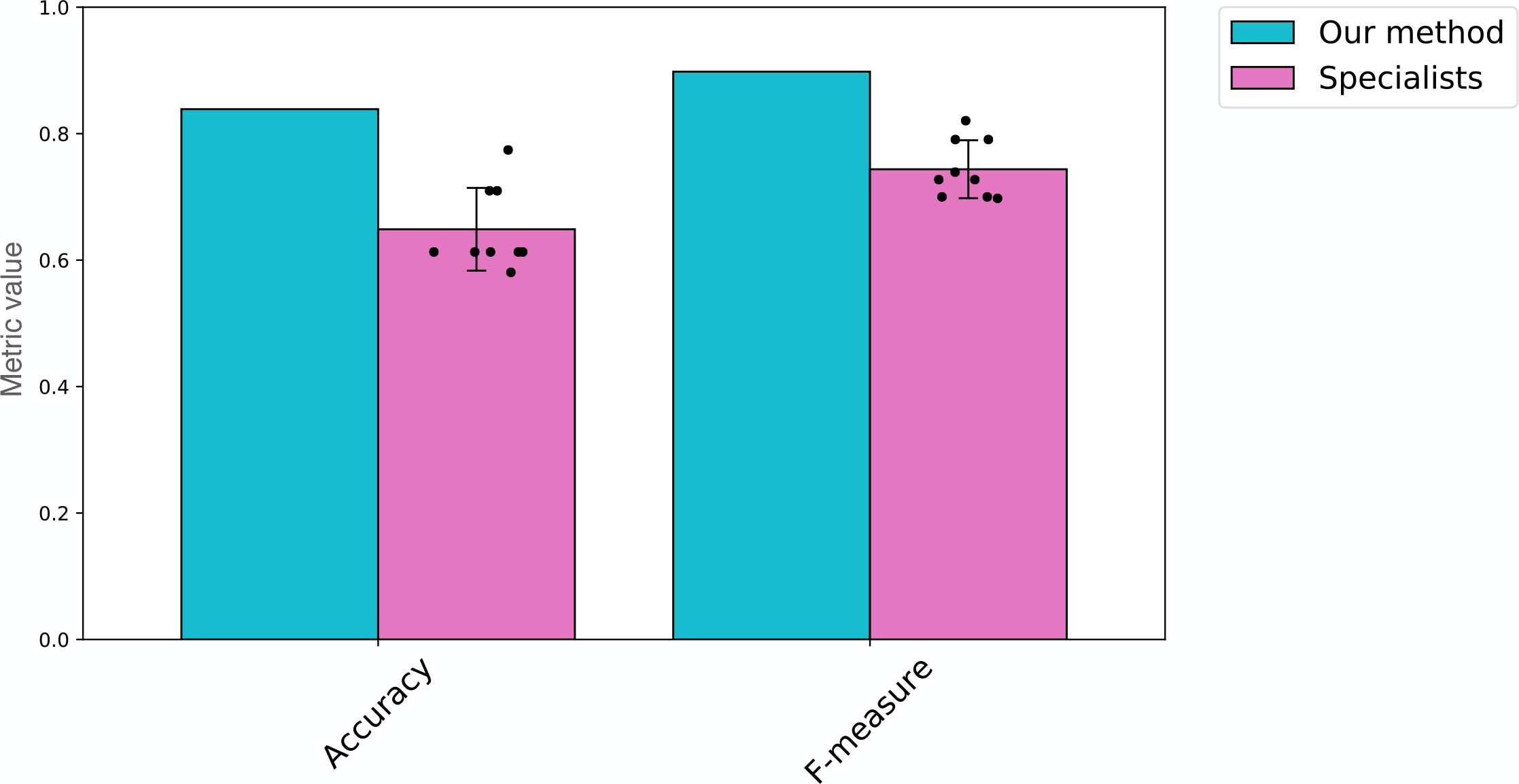
Comparison of embryo classification accuracy between our method and nine embryo culture specialists. The accuracy of the specialists is shown as mean and standard deviation; the dots represent the values for each specialist.

### Analysis of embryo behaviour focused on by NVAN when predicting live birth potential

We considered that determining the variables on which NVAN focused in the input time-series data when classifying embryos might allow us to interpret the classification results in terms of embryo behaviour. Therefore, we visualized an “attention map”—a heat map of the degree of attention afforded to each of the 11 variables in the time-series data—for each embryo that NVAN predicted correctly (Supplementary Fig. 2). To further narrow down the points of interest in these attention maps, we calculated and visualized the geometric mean of all the attention maps (Fig. 5a) for each variable at each time point. The results indicated that the classification algorithm paid most attention to the cell nuclei surface area variables, surface_mean and surface_sd, and the cell nuclei aspect variable, aspect_ratio_sd, at the morula stage from 2.0 to 2.5 days post-fertilization.

**Figure 5:**
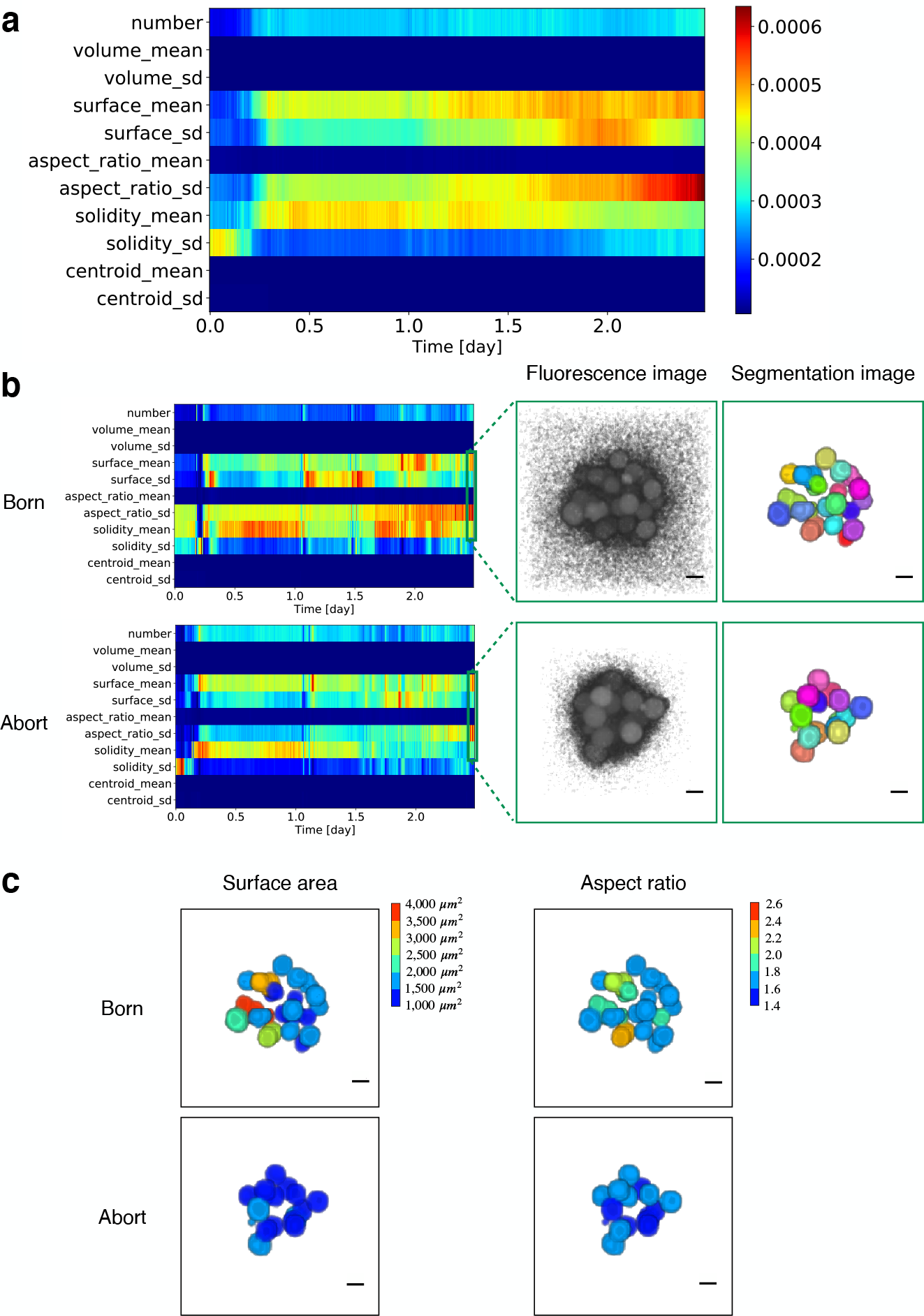
Features that contributed to prediction of the birth potential of all embryos. (a) Heat map of the geometric mean of the attention afforded to each variable and each time point in the embryos that could be correctly classified as born or abort by NVAN. In other words, this heat map showed the features that were common to all embryos in predicting birth embryos. Abbreviations for each variable are as defined in the Figure 2 legend. Warmer colours represent a higher degree of attention and colder colours represent a lower degree of attention. (b) Comparison of the segmentation results for born and abort embryos at the time when the most attention was paid to the surface_mean and aspect_ratio_sd. Heat maps in the left panels show the level of attention given to the indicated variables in the born and abort embryos that were correctly predicted by NVAN. The right panels show the fluorescence images and the segmentation images of cell nuclei. The different colours in the segmentation images indicate the different nuclei. Scale bar, 10 *µm*. (c) Visualization of the surface_mean, surface_sd, and aspect_ratio_sd of each cell nucleus in the segmentation results in panel (b), colour coded by magnitude. Scale bar, 10 *µm*.

To visually understand how these features of interest were differed quantitatively depending on whether the embryo developed to live birth or not, we compared the images of cell nucleus segmentation in born and abort embryos at approximately 2.5 days (Fig. 5b, right), which was when the degree of attention given to surface_mean and aspect_ratio_sd was highest in the NVAN classification of born/abort embryos (Fig. 5b, left). We observed that the cell nuclei were aggregated in both the born and abort embryos at 2.5 days (Fig. 5b, right), suggesting that compaction [30], which is a morphological change in which the cells in the morula stage aggregate and their cell surfaces adhere to each other, might be particularly important for dictating the success or otherwise of the embryo. Furthermore, we visualized the magnitude of the surface_mean, surface_sd, and aspect_ratio_sd of the cell nuclei by colour coding (Fig. 5c). We confirmed that the surface_mean, surface_sd, and aspect_ratio_sd were much larger in the born embryos than in the abort embryos at 2.5 days. We consider that high levels for these variables could potentially be used in combination as a condition for predicting the live birth potential of embryos.

In addition, we determined that the attention given to some variables periodically increased over time in the attention maps of individual embryos (Supplementary Fig. 2). These variables were all related to the shape of the cell nuclei: i.e., surface_mean, surface_sd, aspect_ratio_sd, and solidity_mean. We confirmed that this periodicity coincided with the time at which the cell nuclei increased in number, i.e., immediately after cell division (Fig. 6a). Therefore, we compared the values of these variables in the born embryo group and the abort embryo group immediately after cell division (1st, 2nd, 3rd, 4th mitosis, Fig. 6b). We observed a significant difference between surface_mean immediately after the 4th mitosis and solidity_mean immediately after the 3rd mitosis in the born and abort embryo group (two-sided Wilcoxon rank-sum test, Benjamini-Hochberg adjusted *p*-value = 0.0020, 0.0001 *<* 0.05): surface_mean of the cell nuclei immediately after the 4th mitosis was larger in the born embryo than in the abort embryo, and solidity_mean of cell nuclei immediately after the 3rd mitosis was smaller in the born embryo than in the abort embryo.

**Figure 6:**
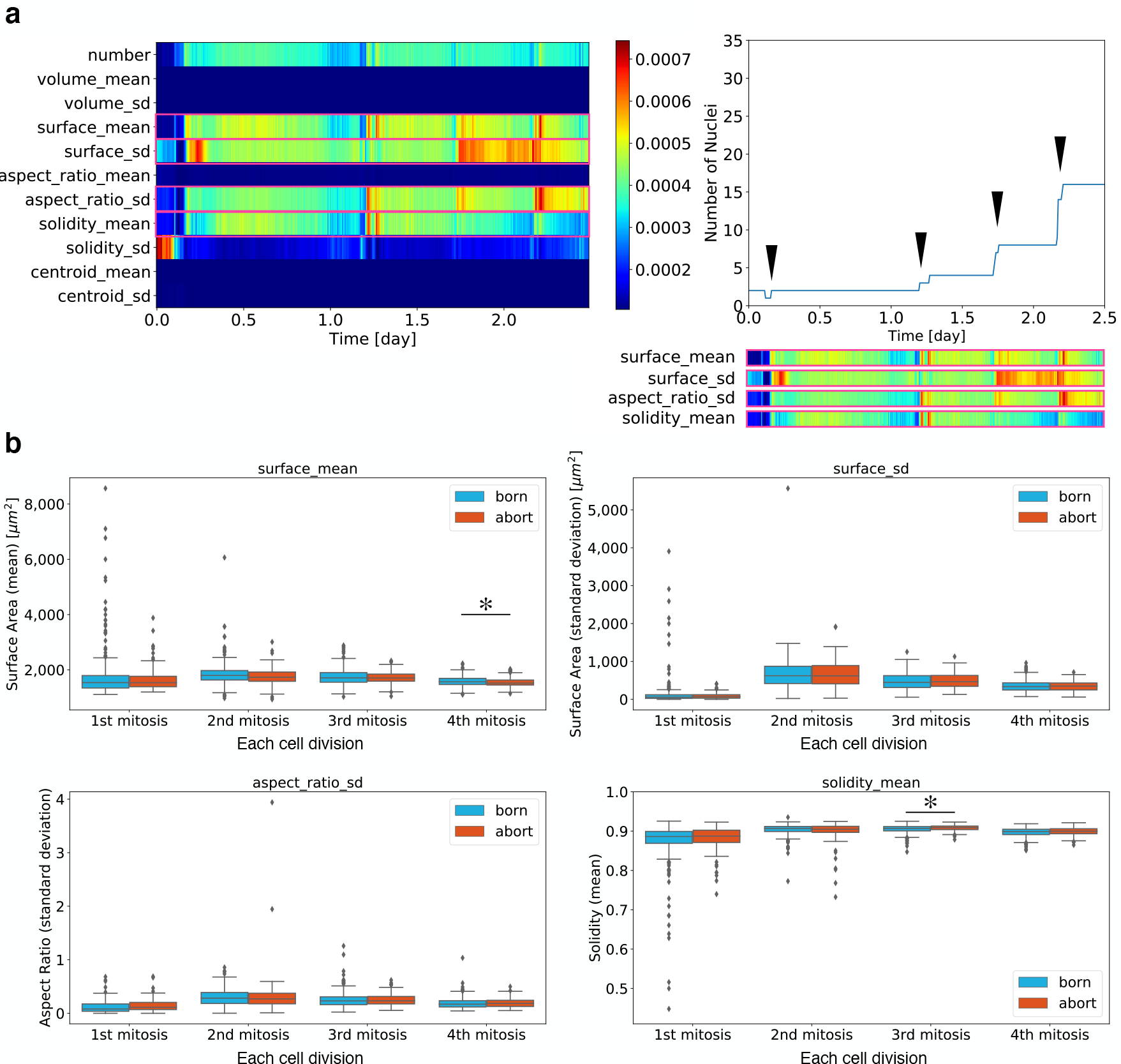
Analysis of characteristic attention patterns in individual embryos. (a) Comparison of the attention heat map of a representative embryo correctly predicted by NVAN and the temporal changes in the number of cell nuclei in that embryo. The timing of the time-periodic increase in the attention given to cell nucleus shape (surface_mean, surface_sd, aspect_ratio_sd, solidity_mean) coincided with the timing of the increase in the number of nuclei due to cell division (green arrowheads). Abbreviations for each variable are as defined in the Figure 2 legend. (b) Box plots of surface_mean, surface_sd, aspect_ratio_sd, and solidity_mean immediately after the 1st mitosis (*n* = 201), 2nd mitosis (*n* = 330), 3rd mitosis (*n* = 603), and 4th mitosis (*n* = 1,170) in the born embryo group and immediately after 1st mitosis (*n* = 93), 2nd mitosis (*n* = 141), 3rd mitosis (*n* = 264), and 4th mitosis (*n* = 489) in the abort embryo group. The bottom and top of the box correspond to the 25th and 75th percentiles, respectively, and the centre line is the median. Whiskers span from the lowest value that is not less than the 25th percentile minus 1.5-fold the interquartile range to the largest value that is not larger than the 75th percentile + 1.5-fold the interquartile range. Data points falling outside this range represent outliers and are reported as dots.∗, Benjamini-Hochberg adjusted *p*-value *<* 0.05, two-sided Wilcoxon rank-sum test.

In summary, we showed that the embryonic behaviours that were important for predicting the live birth potential of mouse embryos were the mean and standard deviation of cell nuclei surface area in the morula stage, the standard deviation of cell nuclei aspect ratio in the morula stage, and the shape of the cell nuclei immediately after cell division. In addition, the results of the attention analysis suggested that, to predict the live birth potential of embryos, NVAN primarily used the criteria that: (a) the mean and standard deviation of surface area and aspect ratio of the cell nuclei at the morula stage are large, (b) the cell nuclei surface area immediately after the 4th mitosis is large, and (c) the cell nuclei solidity immediately after the 3rd mitosis is small. These features might be promising criteria for embryo grading to improve fertility.

## Discussions

We used QCANet to perform image segmentation of cell nuclei in published images of mouse embryos at various stages of embryogenesis, and we developed an algorithm, NVAN, to analyse the resultant multivariate time-series data in order to predict the live birth potential of the embryos. We demonstrated that the embryo classification accuracy of our method was the highest among existing classification methods (Fig. 3).

The classification accuracy with NVAN (Accuracy, 0.8387; F-measure, 0.8980; AUROC, 0.7879; AUPR, 0.8855) was superior to that with MuVAN (Accuracy, 0.7097; F-measure, 0.8235; AUROC, 0.6364; AUPR, 0.7937), which is known to have a high classification accuracy and is a similar algorithm to NVAN. Both these methods have an attention mechanism for focusing on multivariate time-series data, but they use different normalization methods when creating an attention map. Since NVAN performs min-max normalization in the time direction, the difference between the minimum and maximum attention levels became clear. On the other hand, MuVAN uses the softmax function for normalization, so the total attention value in the time direction was 1, meaning that the difference between the minimum and maximum values was occasionally unclear. Owing to these differences, the attention maps generated by NVAN and MuVAN were profoundly different (Supplementary Fig. 3). In the attention map generated by MuVAN, the amount of attention paid to each variable was almost constant in the time direction even though the input was a time series of several hundred time points. We therefore consider that the classification accuracy of MuVAN was lower than that of NVAN because MuVAN had difficulty utilizing features in the time direction.

The direct analysis of mouse images using CLDNN (2D, 3D, or 5D) showed lower classification accuracy than the use of NVAN on extracted multivariate time-series data from the same images (Fig. 3). Previous studies have reported that it is difficult to predict the birth and implantation of human embryos from bright-field images by using a deep learning-based image classification algorithm similar to CLDNN [1, 31]. The reason for the dramatic improvement in the accuracy of prediction by our method compared with that from bright-field images was that our method was able to use not only the shape of embryos but also the 3D shape (e.g, volume and surface area) of nuclei and chromosomes. Because mouse and human embryos undergo similar early developmental processes [32, 33], our result suggests that the prediction accuracy of birth and implantation of human embryos could potentially be improved by applying a similar method to our method to human embryos.

The classification accuracy of 2DCLDNN exceeded that of embryo culture specialists (Figs. 3, 4) using the same time-series of bright-field microscopic images. This suggests that classification of embryos was a difficult task for humans, and the basis for their judgments was not well defined. Our method, which exceeded the accuracy of 2DCLDNN, which in turn exceeded the accuracy of classification by specialists, therefore has the potential to become a basic technology in the field of developmental engineering.

An interesting question is how NVAN achieved high accuracy despite the large variability in the timeseries data of embryo behaviour used as the input (Fig. 2). In previous studies, we found that inter-individual biological variation in developmental processes was responsible for the variability in the quantitative variables representing embryo behaviour [17]. Therefore, we consider that the variability of the time-series data extracted in this study also reflects biological variability. It seems probable, therefore, that NVAN focused on common features that were not buried in biological variability to predict the live birth potential of the embryos.

Another possible source of variability in the time-series data representing embryo behaviour was noise due to QCANet segmentation error. QCANet is currently the most accurate cell nucleus segmentation algorithm for early mouse embryos [17]. We have previously reported that the accuracy of QCANet is high up to 2.5 days post-fertilization [17], and time-series data after this point were excluded from the data used in this study. Therefore, we consider that the influence of the segmentation error on variation in the time-series data used for training of NVAN was very small.

To explore how noise caused by segmentation errors could affect the learning of NVAN, we compared the classification accuracy achieved by training NVAN with data up to about 3.5 days post-fertilization with that achieved with data up to 2.5 days. The accuracy of NVAN was greatly reduced when the data set was extended to 3.5 days (Supplementary Fig. 4), indicating that the segmentation errors from QCANet might adversely affect the NVAN learning for embryo classification. Therefore, improving the segmentation accuracy of QCANet for embryos after 2.5 days post-fertilization could potentially allow our classification method to be extended to later embryos.

Finally, we considered the biological significance of the embryo behaviour that our method focused on when predicting live birth potential. Our results indicate that the live birth potential of an embryo was particularly high when the mean and standard deviation of the cell nucleus surface area and the standard deviation of the cell nucleus aspect ratio were large at the morula stage. Each cell differentiates into either inner cell mass (ICM) or trophoblastoderm (TE) from the morula stage to the blastocyst stage in early development [34]. Previous studies have reported differences in the size and shape of cell nuclei of ICM and TE in blastocysts [35, 36]. We therefore consider it possible that the differences between cell nuclei in the morula stage on which our method focused in the born and abort embryos had already occurred by the time the cells were preparing to differentiate into ICM and TE.

Our results also suggest that the live birth potential of embryos was high when the mean of cell nucleus surface area immediately after 4th mitosis was large and the mean of cell nucleus solidity immediately after 3rd mitosis was small (Fig. 6b). The 3rd mitosis and 4th mitosis are the cell divisions that lead to the morula stage [34, 37]. Previous studies have reported that cell shape during the morula stage differs between inner and outer cells [38], which differentiate into ICM and TE, respectively [39, 40]. Therefore, it is possible that one focus of our method—the shape of the cell nucleus during cell division—represents a feature that determines the cell fate for inducing cell differentiation into ICM and TE.

Another feature to which our method paid attention was the number of cell nuclei during the period between one cell division and the next (Fig. 6a). This feature represented the time of the synchrony of cell division. The synchrony of cell division has been reported to be an important feature of normal embryonic development [17]. The results of our method suggested that the synchrony of cell division was also an important contributor to live birth potential.

The usefulness of our method was confirmed only for mouse embryos. However, we anticipate that it would also have high accuracy when applied to human and bovine embryos and so could be used to help determine the embryogenesis behaviour required for live birth. It should be noted that, to use our method, it is necessary to perform segmentation of fluorescence microscopic images of the early embryos acquired by live-cell imaging. In ART for humans and livestock, it is not acceptable to perform live-cell imaging labelled by fluorescent staining because of potential deleterious effects on the offspring. In recent years, some studies have proposed methods to enable highly accurate image analysis, such as segmentation, for data of various modalities (e.g., brightfield microscope images and fluorescence microscope images) [41–44]. If these methods can be used to perform segmentation of brightfield microscopic images of early embryos in the future, our method could potentially contribute to ART for humans and livestock.

## Methods

### Animals

Some of the live-cell imaging data used in this study were reported previously [21]; new data were obtained under the same permits of the animal committee as described below, under the same rearing environment. This study conformed to the Guide for the Care and Use of Laboratory Animals. All animal experiments were approved by the Animal Care and Use Committee at the Research Institute for Kindai University (permit number: KABT-31–016). ICR or B6D2F1 (BDF1) strain mice (12–16 weeks old) were obtained from Japan SLC, Inc. (Shizuoka, Japan). Room conditions were standardized, with the temperature maintained at 23°C, relative humidity at 50%, and a 12-h/12-h light-dark cycle. Animals had free access to water and commercial food pellets. Mice used for experiments were sacrificed by cervical dislocation.

### The single blastocyst transfer dataset

The single blastocyst transfer dataset reported in [21] is a time series of fluorescence microscopy images of early mouse embryos fluorescently labelled with the histone H2B–mCherry probe. The dataset was obtained by live-cell imaging from immediately after fertilization until 3.5 days post-fertilization. The developmental progress of each embryo was followed, and the embryos were labelled as born (62 embryos) or abort (29 embryos) [21]. The total number of fluorescence microscopic images was 2,359,200. The imaging conditions are shown in Supplementary Table 3.

### Acquiring time series data using QCANet

By using QCANet [17], which has already been trained for segmenting early mouse embryos, we segmented time-series fluorescence microscopic images of the single blastocyst transfer dataset described above. Then, the following 11 variables were measured as time-series data from the time-series segmentation images of each embryo:

- number of cell nuclei (number)
- mean volume of cell nucleus (volume_mean)
- standard deviation of cell nucleus volume (volume_sd)
- mean surface area of cell nucleus (surface_mean)
- standard deviation of cell nucleus surface area (surface_sd)
- mean aspect ratio of cell nucleus (aspect_ratio_mean)
- standard deviation of cell nucleus aspect ratio (aspect_ratio_sd)
- mean solidity of cell nucleus (solidity_mean)
- standard deviation of cell nucleus solidity (solidity_sd)
- mean distance between embryo centre and every cell nucleus (centroid_mean)
- standard deviation of the distance between embryo centre and every cell nucleus (centroid_sd)

### NVAN overview

NVAN is an algorithm that we developed in this work to predict the live birth potential of an embryo from the input of multivariate time-series data representing embryo behaviour (Fig. 1). NVAN consists of four processing compartments (View-wise Recurrent Encoder, Normalized Hybrid Focus, Multi-view Attention, and View-wise Attentional Feature Fusion), as described in the sections below.

#### View-wise Recurrent Encoder

In the input multivariate time-series data, the time is represented as *t* (1, …, *T*), the variable is represented as *v* (1, …, *V*), and the data themselves are expressed as 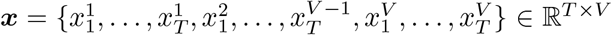. In the processing of View-wise Recurrent Encoder, the time-series data 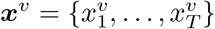 for each variable *v* were input to the Bidirectional Long Short-Term Memory (BLSTM) [45] to acquire the hidden matrix 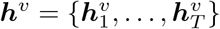, as described by the following formula:

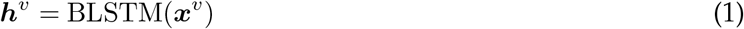

#### Normalized Hybrid Focus

In the process of Normalized Hybrid Focus, the energy matrix ***e*** in order to obtain the attention matrix ***a*** was calculated from the hidden matrix ***h*** acquired by the View-wise Recurrent Encoder. The energy matrix 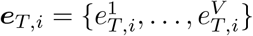 at the time *i* was obtained by Context-based Attention [25], as described by the following formulae:

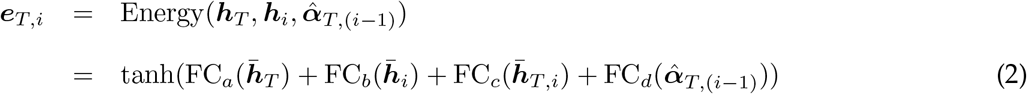

where FC_*a*_, FC_*b*_, FC_*c*_, and FC_*d*_ represent fully connected layers with different weights for 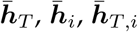, and 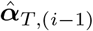, respectively. The context is the relevance and regularity of the subset of data necessary for making predictions in Context-based Attention. 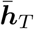 and 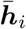 were calculated by the convolution operation to acquire the self-context of the time *T* and attention time *i*, respectively, as follows:

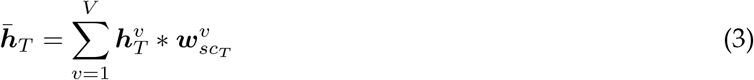

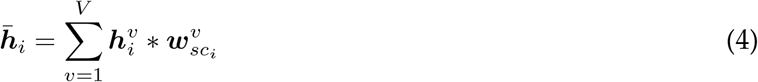

where ∗ represents convolution operator and 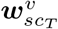 and 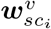 represent the respective convolution kernels. In addition, 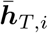 was calculated by the convolution operation to acquire the cross-context of the time *T* and attention time *i*, as follows:

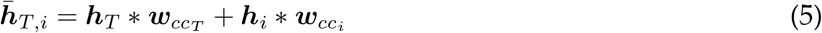

where 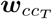 and 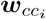 represent the respective convolution kernels. We expected that the relationships between variables would be embedded in the energy matrix ***e***_*T,i*_ by calculating these contexts. Further, 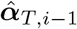 was calculated by the hidden matrix ***h*** at the time *T* and *i −* 1, as follows:

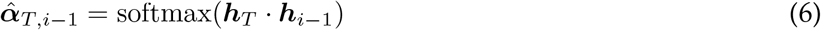

By including 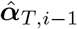 in the calculation for the energy matrix ***e***_*T,i*_, we expected that the relationship between time *i −* 1 and *i* would be embedded in this matrix. Thus, using the above calculation, we were able to obtain the energy matrix ***e***_*T,i*_ considering the relationship between the variables and time. To obtain the attention matrix 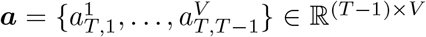 in the multivariate time-series data from the acquired energy matrix ***e***_*T,i*_, normalization was performed as follows:

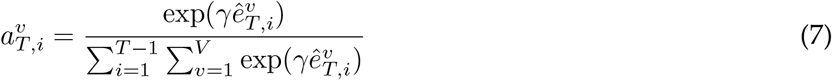

where *γ* represents a sharpening factor [46], which is a constant parameter for controlling the difference between the minimum value and the maximum value of the elements in the normalized matrix of interest. In this method, *γ* = 2. The normalized energy matrix 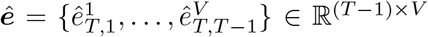 was calculated according to the following equation:

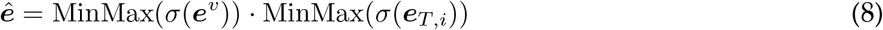

where MinMax represents the min-max normalization function that normalizes to [0, 1] using the minimum and maximum values, and *σ* represents the sigmoid function.

#### Multi-view Attention

In the process of Multi-view Attention, by calculating the context matrix ***c*** from the attention matrix 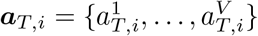 and the hidden matrix 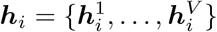, the variate and time context of the hidden matrix was able to be extracted. The context matrix ***c*** is calculated according to the following equation.

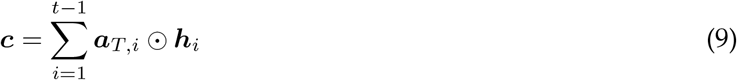

#### View-wise Attentional Feature Fusion

In the process of View-wise Attentional Feature Fusion, 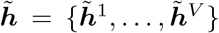 is obtained by applying the convolution operation for each variable *v* to the matrix that combines the hidden matrix 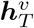 and the context matrix ***c***^*v*^. Furthermore, by inputting 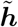 into the fully connected layer, the output ***ŷ*** was obtained as follows:

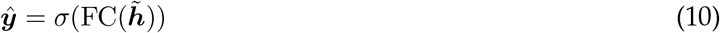

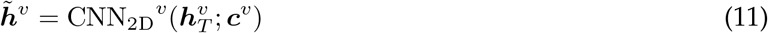

where CNN_2D_^*v*^ represents the convolutional layer for the variable *v*. By performing the convolution operation for each variable in this way, we expected that the born/abort classification of the embryos would be predicted on the basis of the relationships between the variables and the time, which do not depend on the arrangement of the variables in the matrix 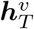 and ***c***^*v*^.

### Training procedure for NVAN

Since NVAN aims to classify embryos as born/abort, the objective function was the binary cross-entropy function for learning binary classification *L*, expressed by the following equation:

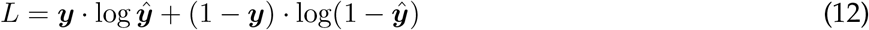

where ***ŷ*** represents the output and ***y*** represents the correct answer. The hyperparameters used in the training of NVANs are shown in Supplementary Table 4. The hyperparameters were determined by using Optuna [47], which can perform Bayesian optimization. To avoid label imbalance problems in the dataset during training, under-sampling [48] was used. The input data (i.e., multivariate time-series data) were subjected to *z*-score normalization, which normalized the mean to 0 and the variance to 1 for each variable, as a pre-process. In addition, as an augmentation of the input data, we removed the input data for the number of time points determined by a uniform random number from the end of the time-series data; this number was updated every epoch (i.e., every pass of the training set completed by the algorithm). By this augmentation, we expected that the learning would be robust to differences in the length of the time-series data.

### Training procedure for other machine-learning methods

The hyperparameter values used for LSTM, AttentionLSTM, and Transformer are shown in Supplementary Tables 5 and 6. The hyperparameter values used for MuVAN, TG-LSTM, and TapNet, which are other classification methods for multivariate time-series data, are shown in Supplementary Tables 7, 8, and 9, respectively. The hyperparameter values used for XGBoost, which is a decision tree method, are shown in Supplementary Table 10. The hyperparameters values used for CLDNN, which was adopted as a method of classifying directly from time-series images, are shown in Supplementary Table 11. All of these learning models were trained under the same conditions as for the NVAN learning method.

### Evaluation procedure for the trained model

The training was evaluated by 4-fold cross-validation by first dividing the dataset into training and test sets at a ratio of 2 : 1 and then further dividing the training data into training and validation sets at a ratio of 3 : 1. The model showing the highest F-measure for the validation data was selected out of all epochs for each fold. The final selected model was evaluated by using the test data. The evaluation metrics of classification were Accuracy, F-measure, AUROC, and AUPR.

### Classification of embryos by embryo cluture specialists

Nine specialists classified the early mouse embryos into born/abort by observing time-series bright-field microscopic images of the single blastocyst transfer dataset. The classified embryos were the same test data created in the above-mentioned division of the dataset. All specialists classified embryos under the condition that they knew the born/abort ratio in the dataset.

### Analysis of important features identified by NVAN in the classification of embryos

Nucleus shapes (surface_mean, surface_sd, aspect_ratio_sd, solidity_mean) immediately after cell division were obtained at 1st, 2nd, 3rd, and 4th mitosis, respectively. The 1st mitosis was the timing when the number of cell nuclei increased between the 1 and 2 cell stages, the 2nd mitosis was the timing when the number of cell nuclei increased between the 2 and 4 cell stages, the 3rd mitosis was the timing when the number of cell nuclei increased between the 4 and 8 cell stages, and the 4th mitosis was the timing when the number of cell nuclei increased between the 8 and 16 cell stages. Nucleus shape was taken from the cell nucleus segmentation image obtained by QCANet at a time within 30 min from these timing.

### Statistical analysis

All statistical analyses in this study were performed by using the two-sided Wilcoxon rank-sum test in SciPy in Python library. The adjusted *p*-value was calculated by using the Benjamini-Hochberg method to correct for multiple testing, and the results were judged to be significant under significance level *α* = 0.05.

### Reporting summary

Further information on research design is available in the Nature Research Reporting Summary linked to this article.

## Acknowledgement

The research was funded by JSPS KAKENHI Grant Number 19J13189 to Y.T., JSPS KAKENHI Grant Number 20H03244 to A.F., JST CREST Grant Number JPMJCR1927 to T.J.K., and JST CREST Grant Number JPMJCR2011 to A.F. and T.J.K. We are grateful for the manual prediction of live birth embryos by members of the Yamagata lab. We are grateful for editing the manuscript carefully by two native-English-speaking professional editors from ELSS, Inc.

## Author contributions

Y.T., K.Y., and A.F. designed the conceptual idea and the study. Y.T. developed and implemented the algorithm of NVAN. D.M. and Z.I. collected the imaging data. Y.T., T.G.Y., and A.F. wrote the manuscript, with suggestions from the other authors.

## References

[1] Khosravi, P. et al. Deep learning enables robust assessment and selection of human blastocysts after in vitro fertilization. NPJ digital medicine 2, 1–9 (2019).

[2] Manna, C., Nanni, L., Lumini, A. & Pappalardo, S. Artificial intelligence techniques for embryo and oocyte classification. Reproductive biomedicine online 26, 42–49 (2013).

[3] Bó, G. & Mapletoft, R. Evaluation and classification of bovine embryos. Animal Reproduction (AR) 10, 344–348 (2018).

[4] Rocha, J. C. et al. A method based on artificial intelligence to fully automatize the evaluation of bovine blastocyst images. Scientific reports 7, 1–10 (2017).

[5] Araki, M. & Ishii, T. International regulatory landscape and integration of corrective genome editing into in vitro fertilization. Reproductive biology and endocrinology 12, 1–12 (2014).

[6] Ueda, J. et al. Heterochromatin dynamics during the differentiation process revealed by the DNA methylation reporter mouse, MethylRO. Stem Cell Reports 2, 910–924 (2014).

[7] Saiz, I. C. et al. The embryology interest group: updating ASEBIR’s morphological scoring system for early embryos, morulae and blastocysts. Medicina Reproductiva y Embriología Clínica 5, 42–54 (2018).

[8] Bó, G. & Mapletoft, R. Evaluation and classification of bovine embryos. Animal Reproduction (AR) 10, 344–348 (2013).

[9] Veeck, L. L. & Zaninovic, N. An atlas of human blastocysts (CRC Press, 2003).

[10] Gardner, D. K., Lane, M., Stevens, J., Schlenker, T. & Schoolcraft, W. B. Blastocyst score affects implantation and pregnancy outcome: towards a single blastocyst transfer. Fertility and sterility 73, 1155–1158 (2000).

[11] Veeck, L. L. Atlas of the human oocyte and early conceptus, vol. 2 (Williams & Wilkins, 1991).

[12] Tian, Y. et al. Predicting pregnancy rate following multiple embryo transfers using algorithms developed through static image analysis. Reproductive biomedicine online 34, 473–479 (2017).

[13] Paternot, G., Debrock, S., De Neubourg, D., d’Hooghe, T. & Spiessens, C. Semi-automated morphometric analysis of human embryos can reveal correlations between total embryo volume and clinical pregnancy. Human reproduction 28, 627–633 (2013).

[14] Viswanath, P., Weiser, T., Chintala, P., Mandal, S. & Dutta, R. Grading of mammalian cumulus oocyte complexes using machine learning for in vitro embryo culture. In 2016 IEEE-EMBS International Conference on Biomedical and Health Informatics (BHI), 172–175 (IEEE, 2016).

[15] Filho, E. S. et al. A method for semiautomatic grading of human blastocyst microscope images. Human Reproduction 27, 2641–2648 (2012).

[16] Adamson, G. D. et al. International committee for monitoring assisted reproductive technology: world report on assisted reproductive technology, 2011. Fertility and sterility 110, 1067–1080 (2018).

[17] Tokuoka, Y. et al. 3D convolutional neural networks-based segmentation to acquire quantitative criteria of the nucleus during mouse embryogenesis. NPJ systems biology and applications 6, 1–12 (2020).

[18] Wong, C. C. et al. Non-invasive imaging of human embryos before embryonic genome activation predicts development to the blastocyst stage. Nature biotechnology 28, 1115–1121 (2010).

[19] Yamamoto, Y. et al. Automated acquisition of explainable knowledge from unannotated histopathology images. Nature communications 10, 1–9 (2019).

[20] Bjerring, J. C. & Busch, J. Artificial intelligence and patient-centered decision-making. Philosophy & Technology 1–23 (2020).

[21] Mashiko, D. et al. Chromosome segregation error during early cleavage in mouse pre-implantation embryo does not necessarily cause developmental failure after blastocyst stage. Scientific reports 10, 1–10 (2020).

[22] Hochreiter, S. & Schmidhuber, J. Long short-term memory. Neural computation 9, 1735–1780 (1997).

[23] Luong, M.-T., Pham, H. & Manning, C. D. Effective approaches to attention-based neural machine translation. In Proceedings of the 2015 Conference on Empirical Methods in Natural Language Processing, 1412–1421 (Association for Computational Linguistics, 2015).

[24] Vaswani, A. et al. Attention is all you need. In Proceedings of the 31st International Conference on Neural Information Processing Systems, 6000–6010 (Curran Associates, Inc., 2017).

[25] Yuan, Y. et al. MuVAN: A multi-view attention network for multivariate temporal data. In 2018 IEEE International Conference on Data Mining (ICDM), 717–726 (IEEE, 2018).

[26] Hu, J. & Zheng, W. Multistage attention network for multivariate time series prediction. Neurocomputing 383, 122–137 (2020).

[27] Zhang, X., Gao, Y., Lin, J. & Lu, C.-T. TapNet: Multivariate time series classification with attentional prototypical network. In Proceedings of the AAAI Conference on Artificial Intelligence, vol. 34, 6845–6852 (AAAI Press, 2020).

[28] Chen, T. & Guestrin, C. XGBoost: A scalable tree boosting system. In Proceedings of the 22nd acm sigkdd international conference on knowledge discovery and data mining, 785–794 (Association for Computing Machinery, 2016).

[29] Sainath, T. N., Vinyals, O., Senior, A. & Sak, H. Convolutional, long short-term memory, fully connected deep neural networks. In 2015 IEEE international conference on acoustics, speech and signal processing (ICASSP), 4580–4584 (IEEE, 2015).

[30] White, M., Bissiere, S., Alvarez, Y. D. & Plachta, N. Mouse embryo compaction. Current topics in developmental biology 120, 235–258 (2016).

[31] Berntsen, J., Rimestad, J., Lassen, J. T., Tran, D. & Kragh, M. F. Robust and generalizable embryo selection based on artificial intelligence and time-lapse image sequences. arXiv preprint 2103.07262 (2021).

[32] Summers, M. C. & Biggers, J. D. Chemically defined media and the culture of mammalian preimplantation embryos: historical perspective and current issues. Human reproduction update 9, 557–582 (2003).

[33] Quinn, P. & Horstman, F. C. Is the mouse a good model for the human with respect to the development of the preimplantation embryo in vitro? Human reproduction 13, 173–183 (1998).

[34] Chazaud, C. & Yamanaka, Y. Lineage specification in the mouse preimplantation embryo. Development 143, 1063–1074 (2016).

[35] Smith, E. R. et al. Nuclear envelope structural proteins facilitate nuclear shape changes accompanying embryonic differentiation and fidelity of gene expression. BMC cell biology 18, 1–14 (2017).

[36] Aiken, C. E., Swoboda, P. P., Skepper, J. N. & Johnson, M. H. The direct measurement of embryogenic volume and nucleo-cytoplasmic ratio during mouse pre-implantation development. Reproduction 128, 527–535 (2004).

[37] Hiiragi, T., Louvet-Vallée, S., Solter, D. & Maro, B. Embryology: does prepatterning occur in the mouse egg? Nature 442, E3–E4 (2006).

[38] Niwayama, R. et al. A tug-of-war between cell shape and polarity controls division orientation to ensure robust patterning in the mouse blastocyst. Developmental cell 51, 564–574 (2019).

[39] Fleming, T. P. A quantitative analysis of cell allocation to trophectoderm and inner cell mass in the mouse blastocyst. Developmental biology 119, 520–531 (1987).

[40] Handyside, A. Time of commitment of inside cells isolated from preimplantation mouse embryos. Development 45, 37–53 (1978).

[41] Lee, G., Oh, J.-W., Her, N.-G. & Jeong, W.-K. DeepHCS++: Bright-field to fluorescence microscopy image conversion using multi-task learning with adversarial losses for label-free high-content screening. Medical image analysis 70, 101995 (2021).

[42] Tokuoka, Y., Suzuki, S. & Sugawara, Y. An inductive transfer learning approach using cycle-consistent adversarial domain adaptation with application to brain tumor segmentation. In Proceedings of the 2019 6th International Conference on Biomedical and Bioinformatics Engineering, 44–48 (Association for Computing Machinery, 2019).

[43] Zhu, Q., Du, B. & Yan, P. Boundary-weighted domain adaptive neural network for prostate mr image segmentation. IEEE transactions on medical imaging 39, 753–763 (2019).

[44] Zhang, Y., Miao, S., Mansi, T. & Liao, R. Task driven generative modeling for unsupervised domain adaptation: Application to X-ray image segmentation. arXiv preprint 1806.07201 (2018).

[45] Schuster, M. & Paliwal, K. K. Bidirectional recurrent neural networks. IEEE transactions on Signal Processing 45, 2673–2681 (1997).

[46] Chorowski, J., Bahdanau, D., Serdyuk, D., Cho, K. & Bengio, Y. Attention-based models for speech recognition. arXiv preprint 1506.07503 (2015).

[47] Akiba, T., Sano, S., Yanase, T., Ohta, T. & Koyama, M. Optuna: A next-generation hyperparameter optimization framework. In Proceedings of the 25th ACM SIGKDD international conference on knowledge discovery & data mining, 2623–2631 (Association for Computing Machinery, 2019).

[48] Wallace, B. C., Small, K., Brodley, C. E. & Trikalinos, T. A. Class imbalance, redux. In 2011 IEEE 11th international conference on data mining, 754–763 (IEEE, 2011).

