## Supplemental Material for "Deep learning-based algorithm for predicting the live birth potential of mouse embryos"

### Supplementary Figures

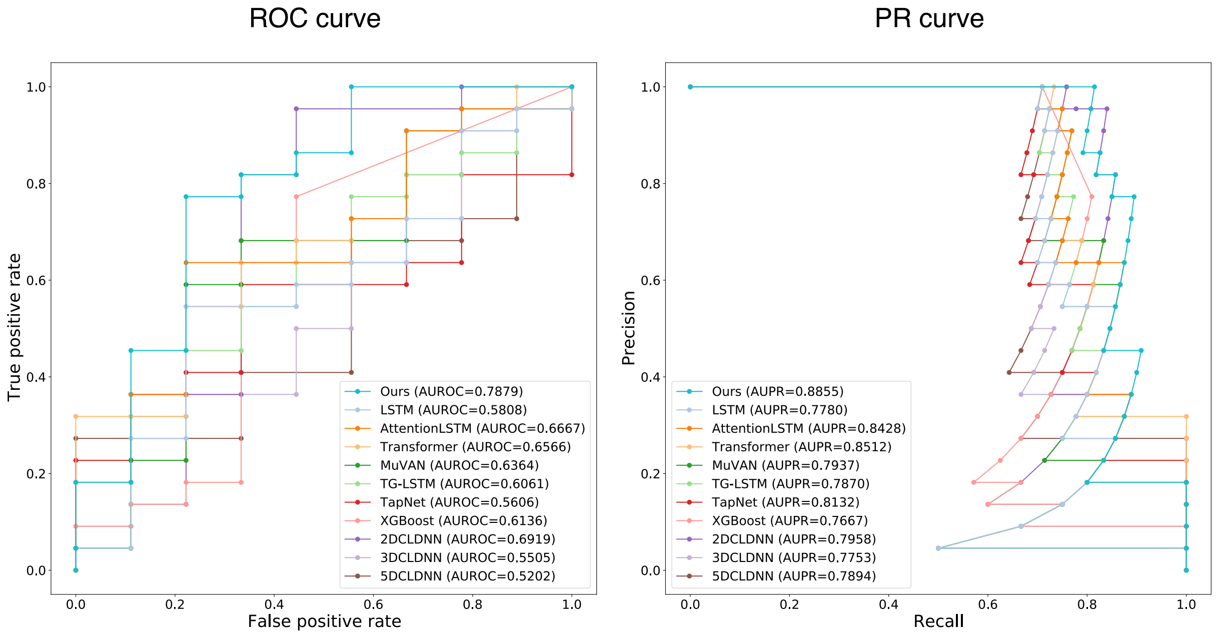


Supplementary Figure 1: **Comparison of the receiver operating characteristic (ROC) curve and precision-recall (PR) curve of each method.** To verify the superiority of the accuracy by our method (Ours), we compared the accuracy with LSTM [1], AttentionLSTM [2], Transformer [3], MuVAN [4], TG-LSTM [5], TapNet [6], XGBoost [7], 2DCLDNN [8], 3DCLDNN [8], 5DCLDNN [8].

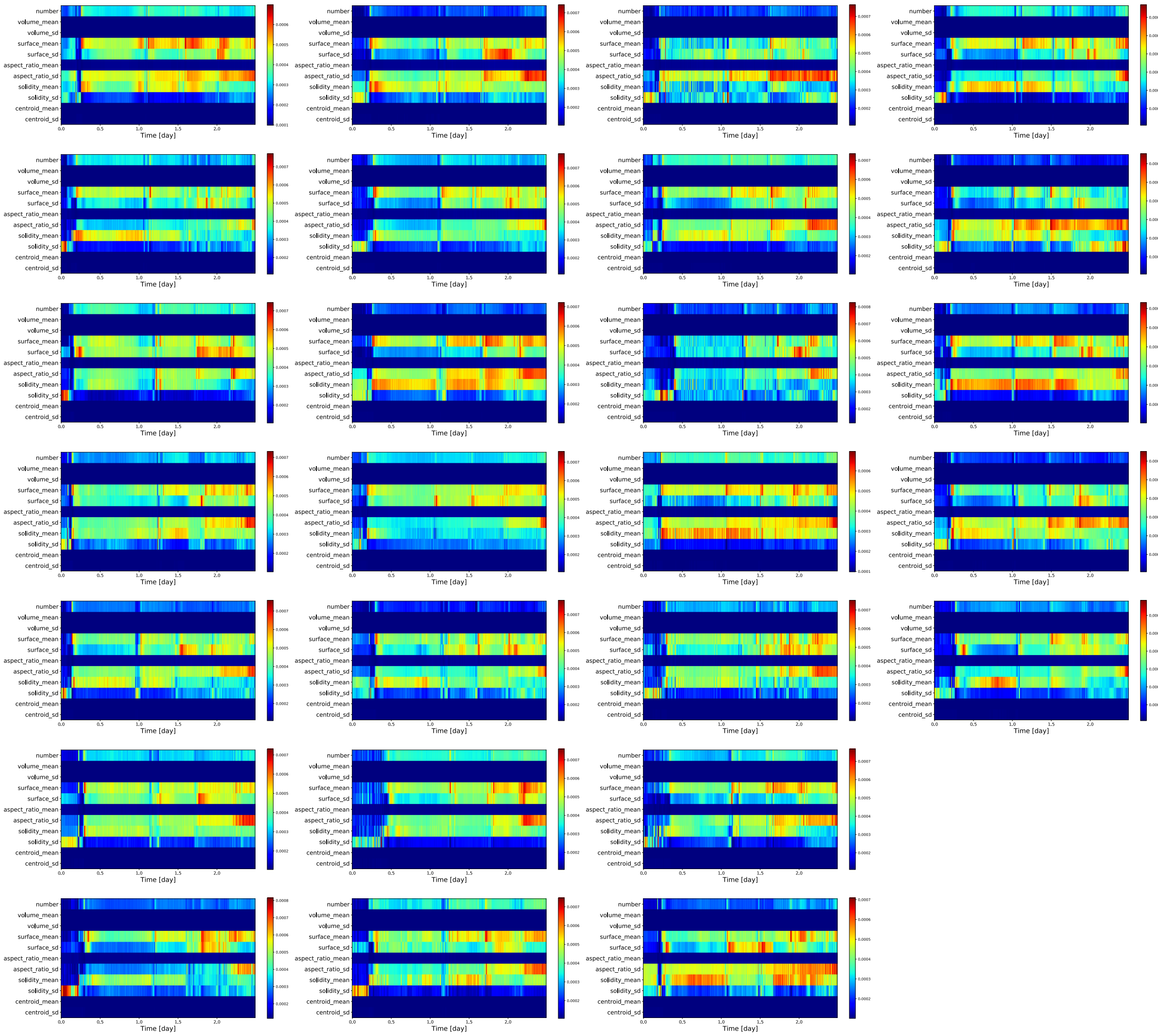


Supplementary Figure 2: **Attention maps of all 26 embryos correctly predicted by Normalized Multi-View Attention Network (NVAN) from the multivariate time-series data.** Such series were number of cell nuclei (number), mean volume of cell nuclei (volume_mean), standard deviation of cell nuclei volume (volume_sd), mean surface area of cell nuclei (surface_mean), standard deviation of cell nuclei surface area (surface_sd), mean aspect ratio (major axis/minor axis) of cell nuclei (aspect_ratio_mean), standard deviation of cell nuclei aspect ratio (aspect_ratio_sd), mean solidity (ratio of total area of the nucleus to the area of the convex hull) of cell nuclei (solidity_mean), standard deviation of cell nuclei solidity (solidity_sd), mean distance between embryo centre and every cell nucleus (centroid_mean), and standard deviation of the distance between embryo centre and every cell nucleus (centroid_sd).

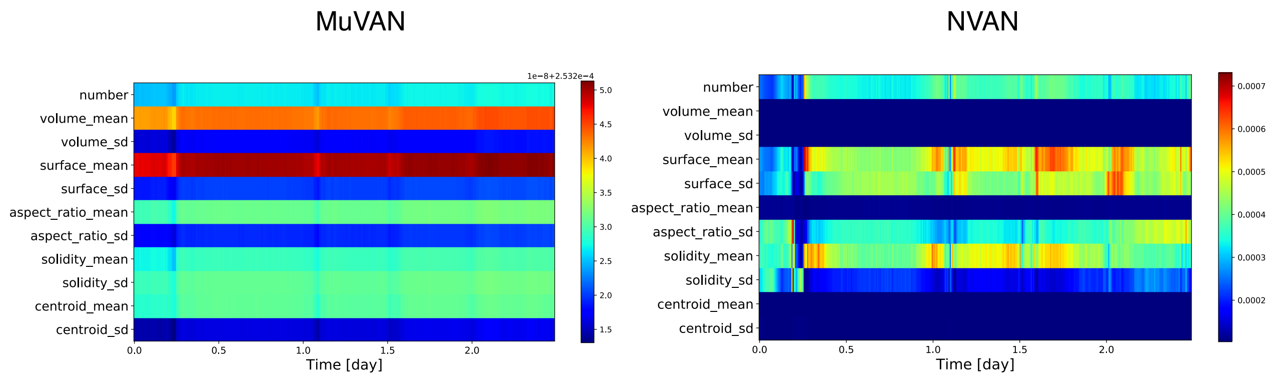


Supplementary Figure 3: **Comparison of the attention maps of NVAN and MuVAN.** These heatmaps represent the attention scores for one representative embryo that was accurately predicted by NVAN and MuVAN [4].

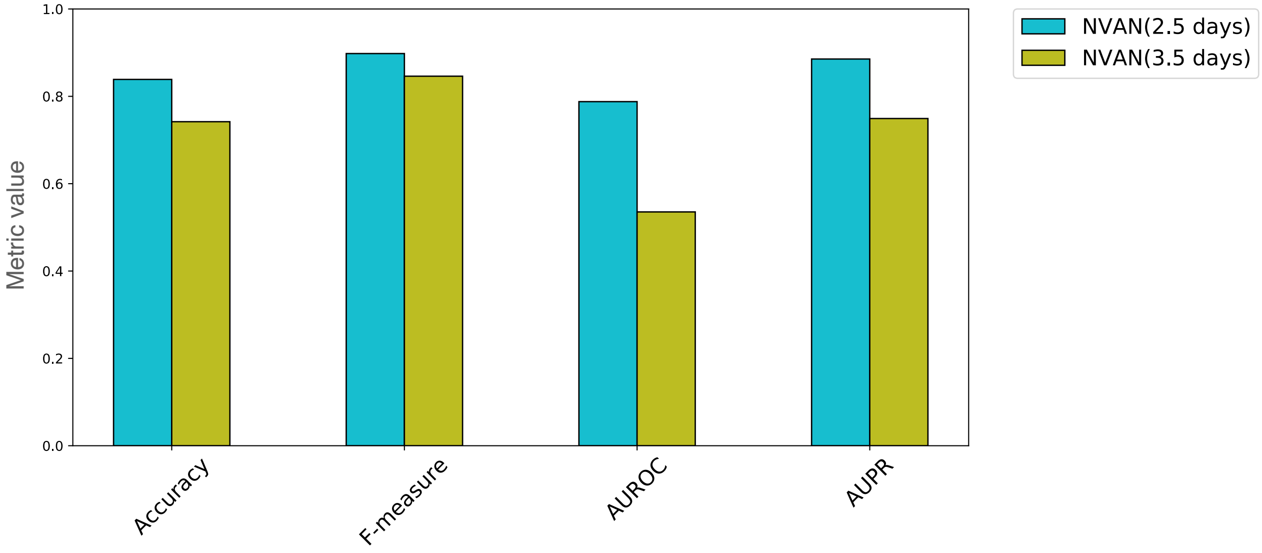


Supplementary Figure 4: **Comparison of the accuracy of NVAN when trained on data up to day 2.5 versus day 3.5 post-fertilization.**

### Supplementary Tables

Supplementary Table 1: Comparison of the classification accuracy of each method. Bold letters indicate the highest metric values among the methods.

| Method | Accuracy | F-measure | AUROC | AUPR |
|---|---|---|---|---|
| Ours | **0.8387** | **0.8980** | **0.7879** | **0.8855** |
| LSTM | 0.6774 | 0.8000 | 0.5808 | 0.7780 |
| AttentionLSTM | 0.6774 | 0.7826 | 0.6667 | 0.8428 |
| Transformer | 0.6452 | 0.7442 | 0.6566 | 0.8512 |
| MuVAN | 0.7097 | 0.8235 | 0.6364 | 0.7937 |
| TG-LSTM | 0.7097 | 0.8302 | 0.6061 | 0.7870 |
| TapNet | 0.6129 | 0.6842 | 0.5606 | 0.8132 |
| XGBoost | 0.6774 | 0.7619 | 0.6136 | 0.7667 |
| 2DCLDNN | 0.7419 | 0.8462 | 0.6919 | 0.7958 |
| 3DCLDNN | 0.5806 | 0.6829 | 0.5505 | 0.7753 |
| 5DCLDNN | 0.7097 | 0.8302 | 0.5202 | 0.7894 |

Supplementary Table 2: Comparison of embryo classification accuracy by our method and embryo culture specialists. Values for specialists are shown as means (standard deviation). Bold letters indicate the highest metric values among the methods.

| Method | Accuracy | F-measure |
|---|---:|---:|
| Ours | **0.8387** | **0.8980** |
| Specialists | 0.6422 (0.0591) | 0.7314 (0.0471) |

Supplementary Table 3: Acquisition conditions of time-series 3D fluorescence microscopic images of mouse embryos for the single blastocyst transfer dataset [9]

| | |
|---|---|
| Observation target | Mouse embryo |
| Fluorescent protein | H2B-mCherry |
| Delivery of fluorescent protein | mRNA Microinjection |
| Microscope and confocal system | CV1000 (Yokogawa Electric Corp, Tokyo, Japan) |
| Image size [$voxel$] | 512×512×51 |
| Spatial resolution ($x : y : z$) [$\mu m/voxel$] | 0.8 : 0.8 : 2.0 |
| Time resolution [$min$] | 10 |
| Number of time slices | 488-525 |
| Number of embryos analysed | 91 |

Supplementary Table 4: Hyperparameters of NVAN

| Hyperparameter | Value |
|---|---|
| Number of LSTM layers | 1 |
| Dimensionality | 128 |
| Dropout | 0.35 |
| Batch size | 4 |
| Optimizer | Adadelta (lr = 0.01) |
| Weight decay | $1.2 \times 10^{-5}$ |
| Epoch | 50 |

Supplementary Table 5: Hyperparameters of LSTM [1] and AttentionLSTM [2]

| Hyperparameter | Value |
|---|---:|
| Number of LSTM layers | 2 |
| Dimensionality | 128 |
| Dropout | 0.5 |
| Batch size | 4 |
| Optimizer | Adadelta (lr = 1.0) |
| Weight decay | $1.0 \times 10^{-3}$ |
| Epoch | 50 |

Supplementary Table 6: Hyperparameters of Transformer [3]

| Hyperparameter | Value |
| --- | ---: |
| Number of self-attention layers | 6 |
| Dimensionality | 256 |
| Dropout | 0.5 |
| Batch size | 4 |
| Optimizer | Adadelta (lr = 1.0) |
| Weight decay | $1.0 \times 10^{-3}$ |
| Epoch | 50 |

Supplementary Table 7: Hyperparameters of MuVAN [4]

| Hyperparameter | Value |
|---|---:|
| Number of LSTM layers | 2 |
| Dimensionality | 128 |
| Dropout | 0.5 |
| Batch size | 4 |
| Optimizer | Adadelta (lr = 1.0) |
| Weight decay | $1.0 \times 10^{-3}$ |
| Epoch | 50 |

Supplementary Table 8: Hyperparameters of TG-LSTM [5]

| Hyperparameter | Value |
|---|---:|
| Number of LSTM layers | 1 |
| Dimensionality | 32 |
| Dropout | 0.0 |
| Batch size | 4 |
| Optimizer | AdaHMG (lr = 0.001) |
| Weight decay | 0.0 |
| Epoch | 100 |

Supplementary Table 9: Hyperparameters of TapNet [6]

| Hyperparameter | Value |
|---|---|
| Number of LSTM layers | 1 |
| Dimensionality | 128 |
| Dropout | 0.0 |
| Batch size | 91 |
| Optimizer | Adam (lr = $1.0 \times 10^{-5}$) |
| Weight decay | $1.0 \times 10^{-3}$ |
| Epoch | 3,000 |

Supplementary Table 10: Hyperparameters of XGBoost [7]

| Hyperparameter | Value |
|---|---|
| Depth | 1 |
| Learning rate | 0.001 |
| Gamma | 0 |
| Estimators | 10,000 |

Supplementary Table 11: Hyperparameters of CLDNN [8]

| Hyperparameter | Value |
|---|---:|
| Kernel size of Convolutional layer 1 | 5 |
| Filter size of Convolutional layer 1 | 8 |
| Kernel size of Pooling layer 1 | 2 |
| Kernel size of Convolutional layer 2 | 5 |
| Filter size of Convolutional layer 2 | 16 |
| Kernel size of Pooling layer 2 | 2 |
| Number of LSTM layers | 2 |
| Dimensionality | 128 |
| Dropout | 0.5 |
| Batch size | 4 |
| Optimizer | Adadelta (lr = 1.0) |
| Weight decay | $1.0 \times 10^{-3}$ |
| Epoch | 50 |

### References


[1] Hochreiter, S. & Schmidhuber, J. Long short-term memory. *Neural computation* **9**, 1735–1780 (1997).

[2] Luong, M.-T., Pham, H. & Manning, C. D. Effective approaches to attention-based neural machine translation. In *Proceedings of the 2015 Conference on Empirical Methods in Natural Language Processing*, 1412–1421 (Association for Computational Linguistics, 2015).

[3] Vaswani, A. *et al.* Attention is all you need. In *Proceedings of the 31st International Conference on Neural Information Processing Systems*, 6000–6010 (Curran Associates, Inc., 2017).

[4] Yuan, Y. *et al.* MuVAN: A multi-view attention network for multivariate temporal data. In *2018 IEEE International Conference on Data Mining (ICDM)*, 717–726 (IEEE, 2018).

[5] Hu, J. & Zheng, W. Multistage attention network for multivariate time series prediction. *Neurocomputing* **383**, 122–137 (2020).

[6] Zhang, X., Gao, Y., Lin, J. & Lu, C.-T. TapNet: Multivariate time series classification with attentional prototypical network. In *Proceedings of the AAAI Conference on Artificial Intelligence*, vol. 34, 6845–6852 (AAAI Press, 2020).

[7] Chen, T. & Guestrin, C. XGBoost: A scalable tree boosting system. In *Proceedings of the 22nd acm sigkdd international conference on knowledge discovery and data mining*, 785–794 (Association for Computing Machinery, 2016).

[8] Sainath, T. N., Vinyals, O., Senior, A. & Sak, H. Convolutional, long short-term memory, fully connected deep neural networks. In *2015 IEEE international conference on acoustics, speech and signal processing (ICASSP)*, 4580–4584 (IEEE, 2015).

[9] Mashiko, D. *et al.* Chromosome segregation error during early cleavage in mouse pre-implantation embryo does not necessarily cause developmental failure after blastocyst stage. *Scientific reports* **10**, 1–10 (2020).
